# The intrinsic predictability of ecological time series and its potential to guide forecasting

**DOI:** 10.1101/350017

**Authors:** Frank Pennekamp, Alison C. Iles, Joshua Garland, Georgina Brennan, Ulrich Brose, Ursula Gaedke, Ute Jacob, Pavel Kratina, Blake Matthews, Stephan Munch, Mark Novak, Gian Marco Palamara, Björn Rall, Benjamin Rosenbaum, Andrea Tabi, Colette Ward, Richard Williams, Hao Ye, Owen Petchey

## Abstract

Successfully predicting the future states of systems that are complex, stochastic and potentially chaotic is a major challenge. Model forecasting error (FE) is the usual measure of success; however model predictions provide no insights into the potential for improvement. In short, the *realized* predictability of a specific model is uninformative about whether the system is inherently predictable or whether the chosen model is a poor match for the system and our observations thereof. Ideally, model proficiency would be judged with respect to the systems’ *intrinsic* predictability – the highest achievable predictability given the degree to which system dynamics are the result of deterministic v. stochastic processes. Intrinsic predictability may be quantified with permutation entropy (PE), a model-free, information-theoretic measure of the complexity of a time series. By means of simulations we show that a correlation exists between estimated PE and FE and show how stochasticity, process error, and chaotic dynamics affect the relationship. This relationship is verified for a dataset of 461 empirical ecological time series. We show how deviations from the expected PE-FE relationship are related to covariates of data quality and the nonlinearity of ecological dynamics.

These results demonstrate a theoretically-grounded basis for a model-free evaluation of a system’s intrinsic predictability. Identifying the gap between the intrinsic and realized predictability of time series will enable researchers to understand whether forecasting proficiency is limited by the quality and quantity of their data or the ability of the chosen forecasting model to explain the data. Intrinsic predictability also provides a model-free baseline of forecasting proficiency against which modeling efforts can be evaluated.

**Glossary:** **Active information**: The amount of information that is available to forecasting models (redundant information minus lost information; Fig. 1).

**Forecasting error (FE)**: A measure of the discrepancy between a model’s forecasts and the observed dynamics of a system. Common measures of forecast error are root mean squared error and mean absolute error.

**Entropy**: Measures the average amount of information in the outcome of a stochastic process.

**Information**: Any entity that provides answers and resolves uncertainty about a process. When information is calculated using logarithms to the base two (i.e. information in bits), it is the minimum number of yes/no questions required, on average, to determine the identity of the symbol (Jost 2006). The information in an observation consists of information inherited from the past (redundant information), and of new information.

**Intrinsic predictability**: the maximum achievable predictability of a system (Beckage et al. 2011).

**Lost information**: The part of the redundant information lost due to measurement or sampling error, or transformations of the data (Fig. 1).

**New information, Shannon entropy rate**: The Shannon entropy rate quantifies the average amount of information per observation in a time series that is unrelated to the past, i.e., the new information (Fig. 1).

**Nonlinearity**: When the deterministic processes governing system dynamics depend on the state of the system.

**Permutation entropy (PE)**: permutation entropy is a measure of the complexity of a time series (Bandt & Pompe, 2002) that is negatively correlated with a system’s predictability (Garland et al. 2015). Permutation entropy quantifies the combined new and lost information. PE is scaled to range between a minimum of 0 and a maximum of 1.

**Realized predictability**: the achieved predictability of a system from a given forecasting model.

**Redundant information**: The information inherited from the past, and thus the maximum amount of information available for use in forecasting (Fig. 1).

**Symbols, words, permutations**: symbols are simply the smallest unit in a formal language such as the letters in the English alphabet i.e., {“A”, “B”,…, “Z”}. In information theory the alphabet is more abstract, such as elements in the set {“up”, “down”} or {“1”, “2”, “3”}. Words, of length *m* refer to concatenations of the symbols (e.g., up-down-down) in a set. Permutations are the possible orderings of symbols in a set. In this manuscript, the words are the permutations that arise from the numerical ordering of *m* data points in a time series.

**Weighted permutation entropy (WPE)**: a modification of permutation entropy (Fadlallah et al., 2013) that distinguishes between small-scale, noise-driven variation and large-scale, system-driven variation by considering the magnitudes of changes in addition to the rank-order patterns of PE.

## Introduction

Understanding and predicting the dynamics of complex systems are central goals for many scientific disciplines (Weigend & Gershenfeld 1993, Hofman et al. 2017). Ecology is no exception as environmental changes across the globe have led to repeated calls to make the field a more predictive science (Clark 2001, Petchey et al. 2015, Dietze et al. 2018, Dietze 2017). One particular focus is *anticipatory predictions*, forecasting probable future states in order to actively inform and guide decisions and policy (Mouquet et al. 2015, Maris et al. 2018). Robust anticipatory predictions require a quantitative framework to assess ecological forecasting and diagnose when and why ecological forecasts succeed or fail.

Forecast performance is measured by **realized predictability** (see glossary), often quantified as the correlation coefficient between observations and predictions, or its complement, **forecasting error (FE)** measures, such as root mean squared error (RMSE). Hence, realized predictability is in part determined by the model used as for any given system, different models will give different levels of realized predictability. Furthermore, it can be unclear, from realized predictability alone, whether the system is stochastic or the model is a poor choice.

By contrast, the **intrinsic predictability** of a system is an absolute measure that represents the highest achievable predictability (Lorenz 1995, Beckage et al. 2011). The intrinsic predictability of a system can be approximated with model-free measure of time series complexity, such as Lyapunov exponents or **permutation entropy** (Boffetta et al. 2002, Bandt & Pompe 2002, Garland et al. 2015). In principle, intrinsic predictability has the potential to indicate whether the model, data, or system are limiting realized predictability. Thus, if we know the intrinsic predictability of a system and its realized predictability under specific models, the difference between the two is indicative of how much predictability can be improved (Beckage et al. 2011).

Here we formalize a conceptual framework connecting intrinsic predictability and realized predictability. Our framework enables comparative investigations into the intrinsic predictability across systems and provides guidance on where and why forecasting is likely to succeed or fail. We use simulations of the logistic map to demonstrate the behaviour of PE in response to time series complexity and the effects of both process and measurement noise. We confirm a general relationship between PE and FE, using a large dataset of empirical time series and demonstrate how the quality, length, and **nonlinearity** in particular of these time series influences the gap between intrinsic and realized predictability and the consequences for forecasting.

## Conceptual framework

The foundation for linking intrinsic and realized predictability lies in information theory and builds on research demonstrating a relationship between PE and FE for complex computer systems (Garland et al. 2015). Information theory was originally developed by Claude Shannon as a mathematical description of communication (Shannon 1948) but has since been applied across many disciplines. In ecology, several information-theoretic methods have proved useful, including the Shannon biodiversity metric in which the probability of **symbol** occurrences (see Box 1) is replaced by the probability of species occurrences (Jost 2006, Sherwin et al. 2017), and the Akaike Information Criterion (Akaike 1974) which is widely used for comparing the performance of alternative models (Burnham & Anderson 2002). Given its importance to our framework, we first provide an introduction to information theory with special attention to applications for ecological time series. Since our goal is to inform where, when and why forecasting succeeds or fails; we then i) describe how **information** may be partitioned into **new and redundant information** based on permutation entropy, ii) demonstrate how redundant information is exploited by different forecasting models, and iii) examine the relationship between permutation entropy and realized predictability and how it can inform forecasting.

### An information-theoretic perspective

A first step towards predicting the future of any system is understanding *if* the observations of that system contain **information** about the future, i.e. does the system have a memory. The total information in each observation can be thought of as a combination of information that came from past states (i.e., **redundant information)** and information that is only available in the present state (i.e., **new information**).

When there is a substantial amount of information transmitted from the past to the present (figure 1 Aiii), the system is said to be highly redundant. In other words, future states depend greatly on the present and past states. In these cases, very little new information is generated during each subsequent observation of the system and the resulting time series is, in theory, highly predictable (has high intrinsic predictability).

**Figure 1A.**
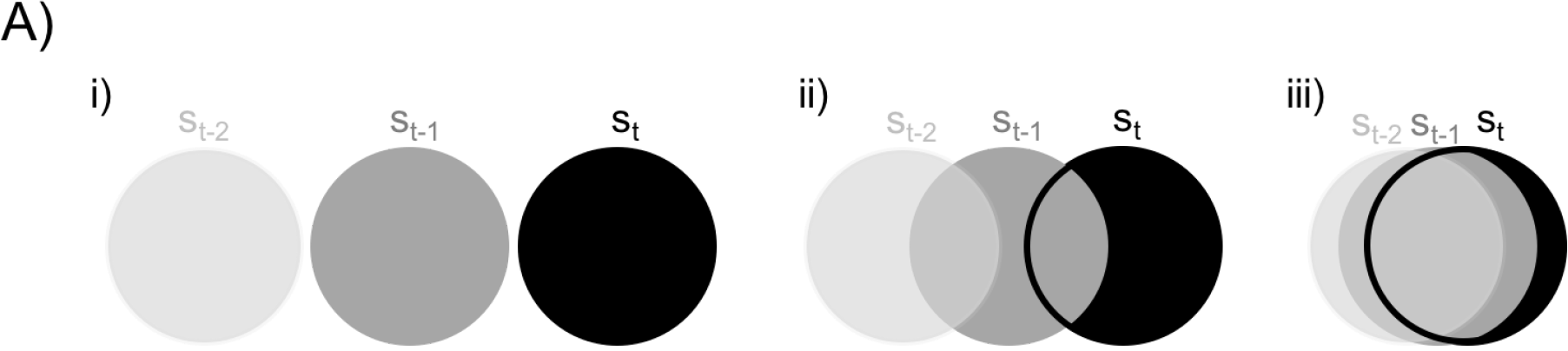
The total information content of an observation of a system at a given state in time, *S*_*t*_, is depicted by filled circles with past states (*S*_*t*−1_ and *S*_*t*−2_) represented by shades of grey, i) lack of overlap between past and present states illustrating a case where no information is transmitted from past states (i.e. a purely stochastic system), with low redundancy and high Shannon entropy rate, ii) intermediate overlap indicating a case when some information is transferred from past to present (i.e. a deterministic system strongly driven by stochastic forcing), with intermediate redundancy and Shannon entropy rate, iii) large overlap indicating a case when the current state is mostly determined by the previous state (i.e. a highly deterministic system), with high redundancy and low Shannon entropy rate. Note that both the redundancy and Shannon entropy rate of a system are intrinsic properties of the system and will only change if the system itself changes.

Conversely, in systems dominated by stochasticity, the system state at each time point is mostly independent of past states (figure 1 Ai). Thus, all of the information will be “new” information, and there will be little to no redundancy with which to train a forecasting model. In this case, regardless of model choice, the system will not be predictable (has low intrinsic predictability).

Imperfect observations introduce uncertainty or bias into time series, and thereby affect the redundant information that is available or perceived. Observation errors in particular will reduce the redundant information available to forecasting models, thus lowering the realized predictability. We refer to this reduction as **lost information,** which is not an innate property of the system but is the result of the practical limitations of making measurements and any information-damaging processing of the data (Fig. 1B, Box 2). As such, lost information can be mitigated and is an important leverage point for ecologists to improve their forecasts. For example, replicate measurements or other forms of data integration that increase estimation accuracy and reduce bias will reduce information loss and can improve forecasts.

**Figure 1B.**
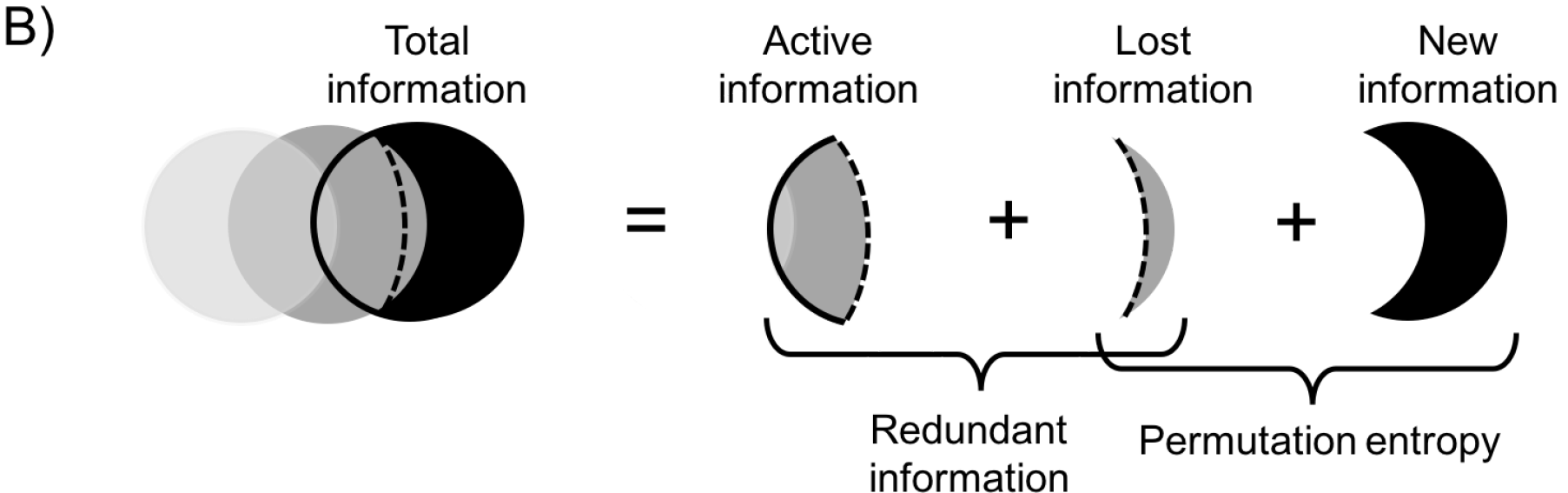
The total information of an observation (black circle) is composed of new information and redundant information; redundant information is composed of active and lost information. A system’s redundancy determines its intrinsic predictability. Information may be lost due to observation error and data processing (*lost information*). This reduces the redundant information that can be used for forecasting (*active information*). Lost information is not an intrinsic property of the system but rather represents practical limitations on our ability to make accurate measurements. The rate at which new information is being generated (Shannon entropy rate) may be approximated with permutation entropy. Because permutation entropy quantifies the joint contribution of the Shannon entropy rate and the lost information, efforts that minimize the amount of lost information not only maximize the redundant information that can actively be used for forecasting but also improve the estimation of the intrinsic Shannon entropy rate.

### Permutation entropy

Permutation entropy (PE) is a measure of time series complexity that approximates the rate at which new information is being generated along a time series (Box 1). PE approximates and is inversely related to intrinsic predictability by quantifying how quickly the system generates new information. Time series with low permutation entropy have high redundancy and are expected to have high intrinsic predictability (Garland et al. 2015).

PE uses a symbolic analysis that translates a time series into a frequency distribution of **words** (see glossary for definition). The frequency distribution of words is then used to assess the predictability of the time series. For example, a time series in which a single word (i.e. a specific pattern) dominates the dynamics has high redundancy and thus future states are well predicted by past states. In contrast, a random time series, in which no single pattern dominates, would produce a nearly uniform frequency distribution of words, with future states occurring independently from past states. Hence, by quantifying the frequency distribution of words, PE approximates how much information is transmitted from the past to the present, corresponding to the intrinsic predictability of a time series.

When observations are imperfect PE measures the joint influence of new information (from either internal or external processes) and lost information (due to the observation process as well as data processing). We refer to the redundant information that is not lost and remains available as **active information,** which is the information that can be exploited by forecasting models.

### Forecasting and redundant information

Realized predictability is highest when the chosen forecasting model exploits all the active information contained in a time series. For illustration, we forecast the oscillating abundance of a laboratory ciliate population (Veilleux 1976) with three different approaches (Figure 1C): i) the mean of the time series (a model which uses relatively little of the active information), ii) a linear autoregressive integrated moving-average model (ARIMA) that uses the local-order structure of the time series in addition to the mean (a model which uses an intermediate amount of the active information), and iii) empirical dynamic modelling (EDM) that can incorporate **nonlinearities,** when present, in addition to the mean and local-order structure (a model which can feasibly use more active information). The time series was split into training data and test data. Forecasting models were fit to the training data and used to make forward predictions among the test data. The forecast performance of the models (i.e. the realized predictability) varied with the amount of information they used, which depended on structural differences among the models that exploit the active information coming from the past. EDM and ARIMA had similar performance suggesting that the time series entailed little nonlinearities for the EDM to exploit.

**Figure 1C.**
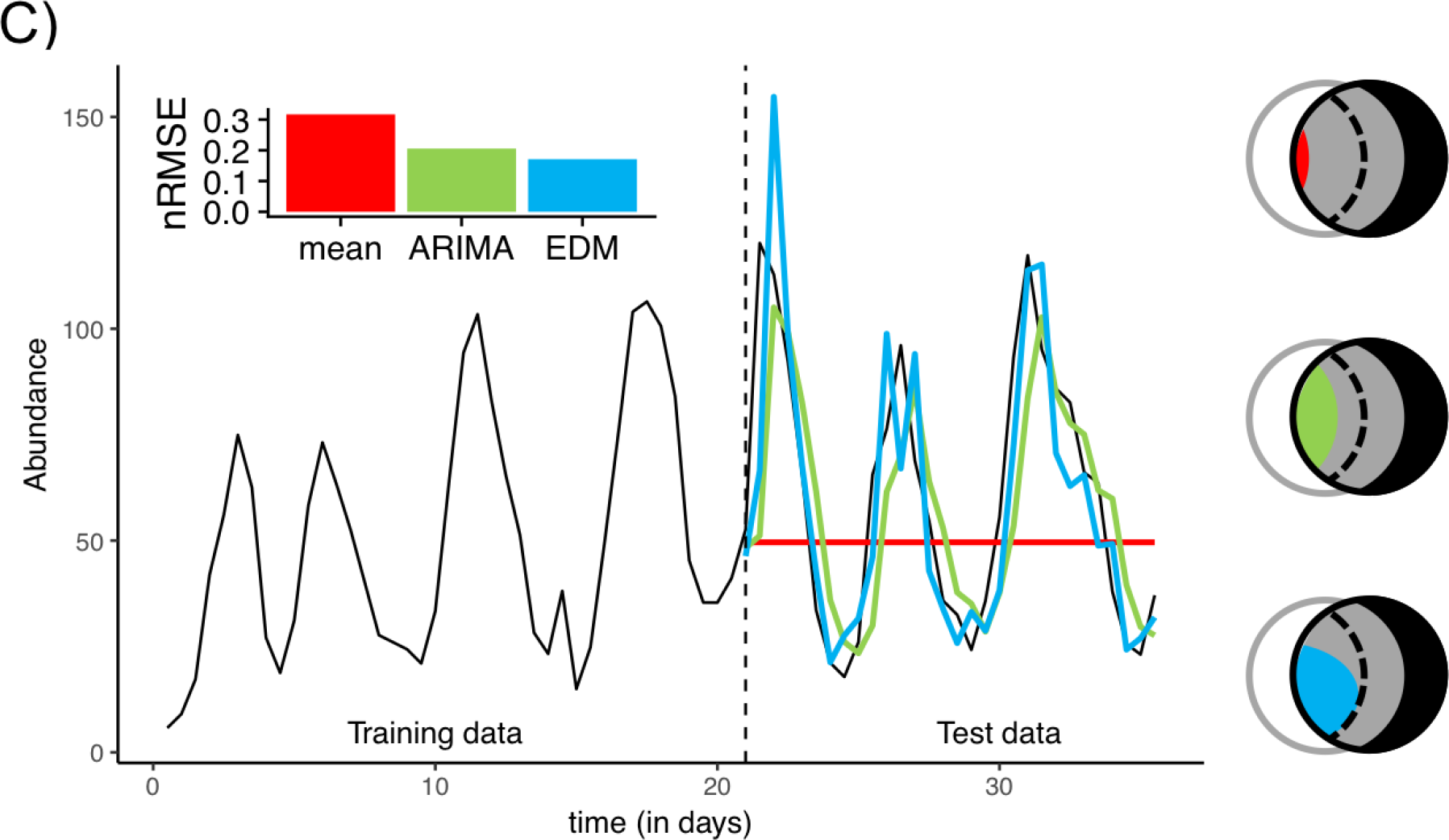
The realized predictability is the degree to which forecast models can exploit the active information of a time series. Consider, for example, a time-series on the abundance of a species (black line) of which the first 21 days are used to train (parameterize) three forecasting models: a forecast that uses the simple mean of the training data set (red), an Autoregressive integrated moving average (ARIMA) model (green), and an Empirical Dynamical Model (EDM, blue). The forecasting performance of these models is assessed using the remaining time series (after day 22). The inset shows the normalized root mean squared error (nRMSE) as a measure of deviation between predicted and observed values (i.e. forecast error) for each of the three forecasting models. ARIMA and EDM exploit the available structure in the data better than the mean forecast, as illustrated by the coloured wedges filling different amounts of the area of active information.

### The relationship between realized and intrinsic predictability

With a perfect forecasting model, realized predictability - measured by forecasting error (FE) - and intrinsic predictability - measured by permutation entropy (PE) - will be positively related. More specifically, the relationship will pass through the origin and monotonically increase up to the maximum limit of PE = 1 (Fig. 1D, the boundary between the white and grey regions; Garland et al. 2015). In the top right of this figure are systems with high PE and therefore low redundancy and high forecasting error. In the bottom-left of the figure are systems with low PE and therefore high redundancy and low forecast error. The boundary is the limit for a perfect model that maximizes the use of active information.

**Figure 1D.**
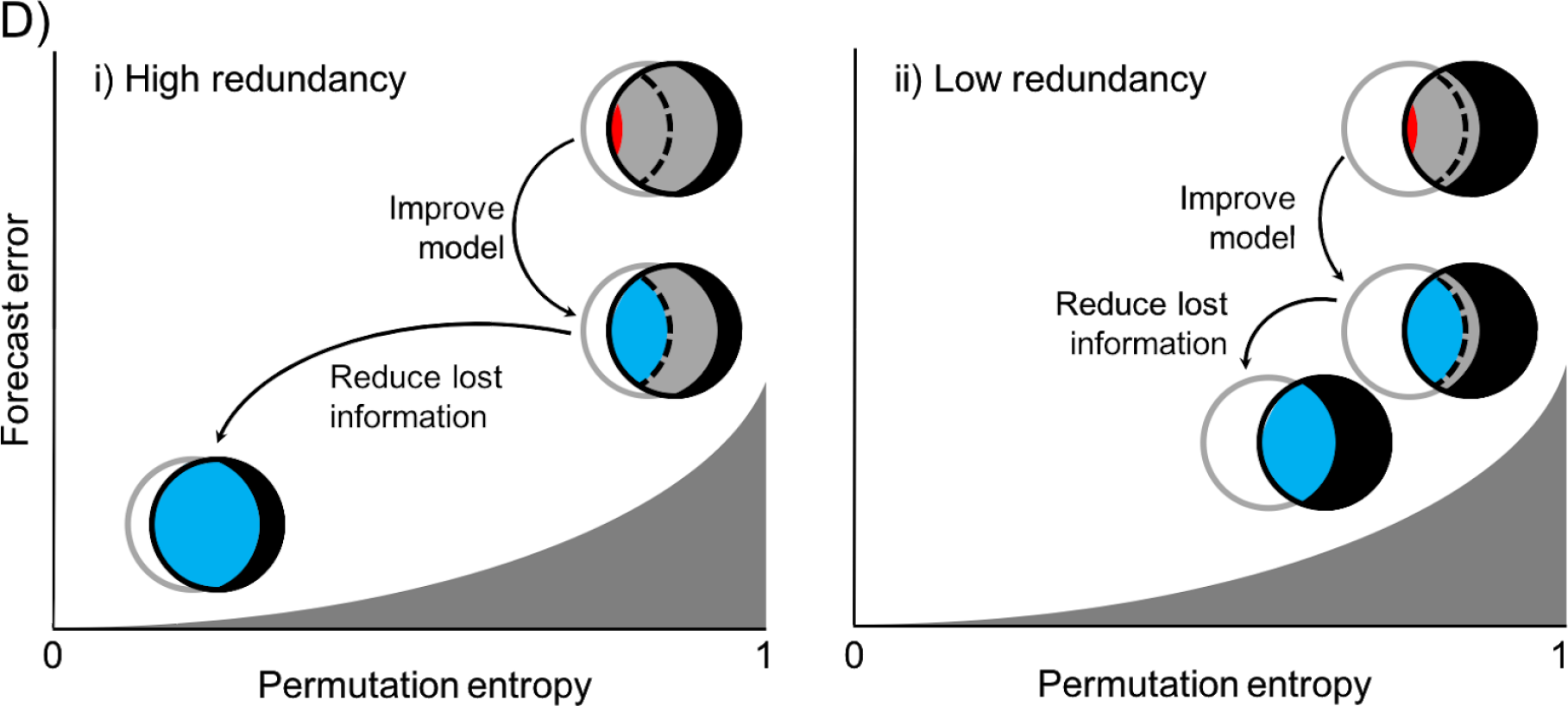
In the relationship between PE and FE a system can be moved toward the ideal grey boundary with forecast models that make better use of active information or by reducing information loss, not necessarily in that order. The two panels depict how to reach the greatest achievable forecasting skill in two different systems that have the same initial permutation entropy but differ in their relative amounts of new and redundant information (i.e. they differ in their intrinsic predictability). As these intrinsic properties of the system cannot be changed, improvements to forecast skill rely on fully exploiting the active information available (e.g., improved forecasting model) and minimizing information loss (e.g., improved sampling) to better approximate the true Shannon entropy rate, which establishes the lower boundary (grey area).

Lost information complicates the interpretation of the PE - FE relationship by obscuring the system’s actual intrinsic predictability. We illustrate this case in figure 1D using two hypothetical systems: one with high intrinsic predictability and a large amount of lost information, and one with lower intrinsic predictability but relatively little lost information. Despite the differences in the redundancy of two systems, the PE of their time series can be very similar (even identical) because PE does not differentiate between new and lost information.

For this example, both systems in figure 1D start with high FE relative to their PE. Selecting more appropriate forecasting models causes a reduction in FE but no change in PE. Reducing lost information (e.g. by increasing the frequency of measurements) decreases both PE and FE. The system with a high redundancy and a low **Shannon entropy rate** has a greater overall potential for improving forecasting skill through the recovery of lost information. In contrast, the system with low redundancy has limited scope to further improve forecasting skill; forecasting is less limited by lost information, but rather by its lower redundancy. As such, the lowest possible forecast error will be substantially higher in the second system than in the first system because the intrinsic predictability of the second is inherently lower and cannot be changed.

### Box 1 Theory and estimation of PE and WPE

Information theory provides several measures for approximating how much new information is expected per observation of a system (e.g. the Shannon-entropy rate and the Kolmogorov-Sinai entropy). However, these measures are only well defined for infinite sequences of discrete random variables and can be quite challenging to approximate for continuous random variables, especially if one only has a finite set of imperfect observations. Permutation entropy is a measure aimed at robustly approximating the Shannon-entropy rate of a times series (or the Kolmogorov-Sinai entropy if the time series is stationary).

Rather than estimating probability mass functions from **symbol** frequencies or frequencies of sequences of symbols, as is done with traditional estimates of the Shannon-entropy rate, permutation entropy uses the frequencies of orderings of sequences of values; it is an ordinal analysis (see Fig. B1-1 below for a visual explanation). The ordinal analysis of a time series maps the successive time-ordered elements of a time series to their value-ordered permutation of the same size. As an example, if [*x*_1_, *x*_2_, *x*_3_] = [11, 6, 8] then its ordinal pattern, or word, ϕ ([*x*_1_, *x*_2_, *x*_3_]), is 2-3-1 since *x*_2_ ≤ *x*_3_ ≤ *x*_1_ (see red time series fragment in box figure I A). PE is calculated by counting the frequencies of these words (or **permutations)** that arise after the time series undergoes this ordinal analysis. That is, given a time series (box figure I A), let *S*_*m*_ be defined as the set of all permutations (possible words) *π* of order (word length) *m* and time delay τ, describing the delay between successive points in the time series (box figure I B for *m* = 3 and τ = 1). For each permutation *π* ∈ *S*_*m*_ we estimate its relative frequency of occurrence for the observed time series after performing ordinal analysis on each delay vector, 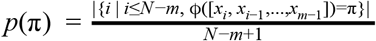, where |·| denotes set cardinality (box figure 1C). Then permutation entropy of order *m* ≥ 2 is calculated as 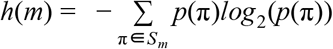.

Since, 0 ≤ *h*(*m*) ≤ *log*_2_(*m*!), it is common to normalize permutation entropy by dividing by *log*_2_(*m*!). With this convention, maximal *h*(*m*) = 1 and minimal *h*(*m*) is equal to 0. Since in the infinite word length limit, permutation entropy is equivalent to the Kolmogorov-Sinai entropy as long as the observational uncertainty is sufficiently small (Amigó et al. 2005), we can approximate the intrinsic predictability of an ecological time series by computing 1 − *h*(*m*).

For the ordinal analysis of a time series, ranks are only well defined if all values are different. If some values are equal (so called ‘ties’), the ordinal analysis is not possible. Several approaches are available to break the ties: the ‘first’ method results in a permutation with increasing values at each index set of ties, and analogously ‘last’ with decreasing values. The ‘random’ method puts these in random order whereas the ‘average’ method replaces them by their mean, and ‘max’ and ‘min’ replaces them by their maximum and minimum respectively, the latter being the typical sports ranking.

In contrast, an ordinal analyses is also affected by small scale fluctuations due to measurement noise which can obscure the influence of large scale system dynamics. **Weighted permutation entropy (WPE)** reduces the influence of small-scale fluctuations by taking into account the relative magnitudes of the time series values within each word (Fadlallah et al. 2013). That is, each word’s (*X*_*t*_ = [*x*_*t*_, *x*_*t*−τ_ …,*x*_*t*−τ(*m* − 1_]) contribution to the probability mass function is weighted by its variance, *viz*., *w*(*X*_*t*_) = *var*(*X*_*t*_). Using this weighting function, the weighted probability of each permutation is estimated by: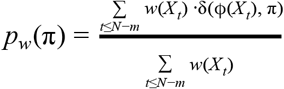 where δ(*x*, *y*) = 1 if and only if *x*=*y* and δ(*x*, *y*) = 0 otherwise. The weighted permutation entropy of order *m* ≥ 2 is then defined as 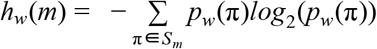. Similar to PE, the weighted permutation entropy is normalized by *log*_2_(*m*!). We use weighted permutation entropy for all analyses presented in this manuscript.

The estimation of PE to time series requires specifying a order *m* and time delay τ. The shorter the chosen word length, the fewer possible words there are and the better we can estimate permutation frequencies. However, the ability to distinguish patterns is limited by the possible number of unique permutations. Hence, when word lengths are too short or too long, the frequency distribution is more uniform. In practice the total length of the time series limits the choice of possible word lengths and hence the number of unique words that can be resolved (Riedl et al. 2013). Regarding the time delay τ, most applications to study the complexity of a time series use a τ = 1 (Riedl et al. 2013). If τ > 1, Bandt (2005) notes the interesting property of the permutation entropy to be small, if the series has main period p for τ = *p*/2 and 3 *p* / 2, and to be large for τ = *p* and τ = 2 *p*. We refer to Riedl et al. 2013 who provide practical considerations regarding setting permutation order *m* and time delay τ.

**Figure B1.**
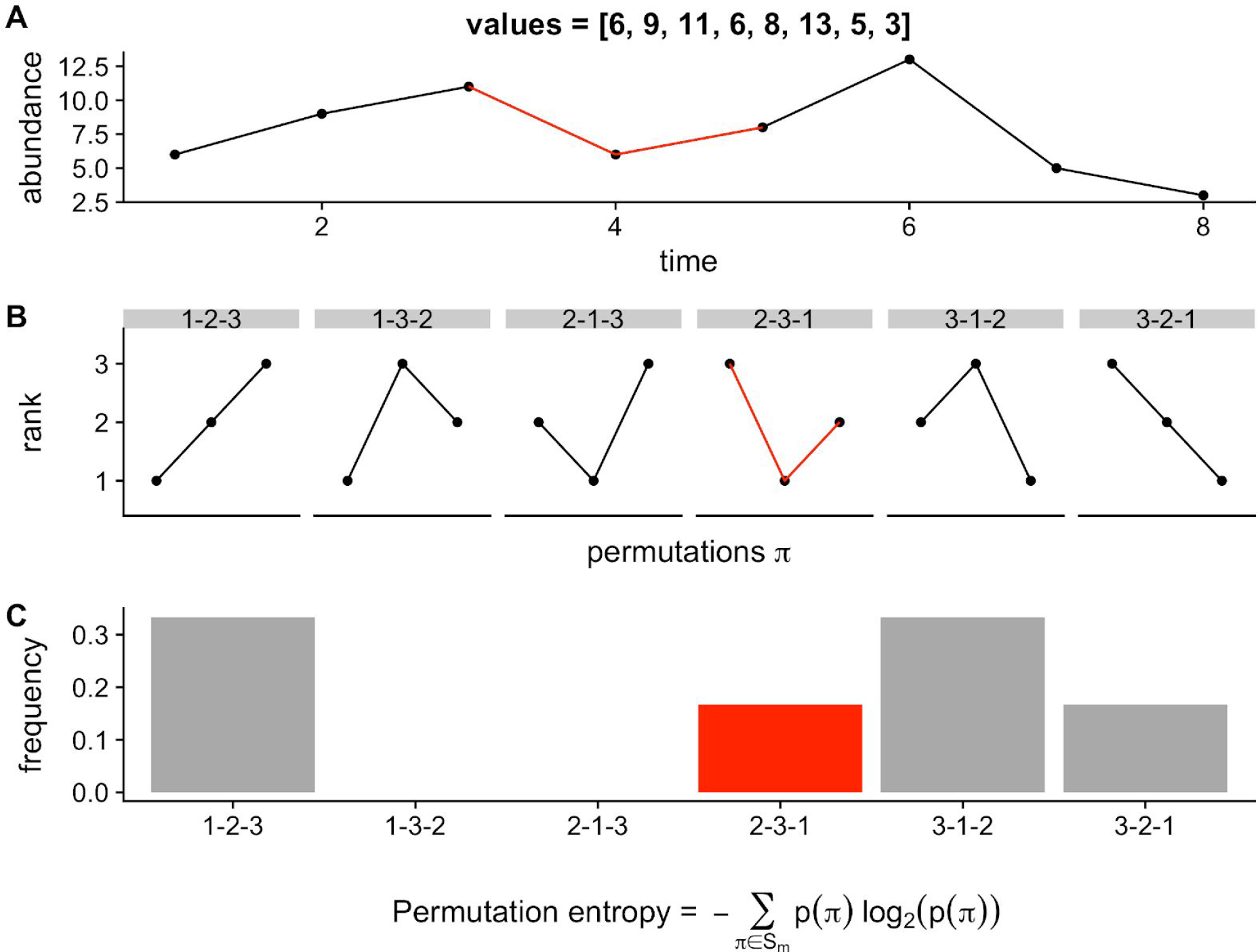
We illustrate how to estimate permutation entropy from an empirical time series (A) assuming m = 3 and τ = 1. A permutation order m = 3 allows for a set of 6 (i.e. 3!) permutations, shown in panel B. The occurrence of each permutation *π* is then counted and divided by the total number of permutations as an estimate of their proportional frequency (panel C). For example, permutations 2-3-1 (shown in red) and 3-2-1 are each only found once in the time series, whereas 1-2-3 and 3-1-2 are found twice, leading to frequencies of 0.17, 0.17, 0.33 and 0.33, respectively. The permutation entropy is then calculated as the Shannon entropy of proportional frequencies. For the given time series this is 1.92, which is normalized by log_2_(3!) yielding a permutation entropy of 0.74.

### Box 2 with information on the limitations of PE/WPE

When analyzing time series, ecologists typically employ a number of data pre-processing methods. These methods are used to reduce low-frequency trends or periodic signals (detrending), reduce high-frequency variation (smoothing), standardize across the time series or reduce the influence of extreme values (transformation), deal with uncertain or missing data points (gap or sequence removal, and interpolation), to examine specific time step sizes (downsampling), or to combine different time series (aggregation). The following table summarizes the anticipated effects on permutation entropy of a suite of commonly used pre-processing methods. In many cases, whether a method increases or decreases permutation entropy will depend on the specific attributes of the time series (e.g., its embedding dimension, autocorrelation, covariance structure, etc.) and the permutation order (*m*) at which its permutation entropy is approximated. This is illustrated by specific examples in Figure B2 which contrasts the permutation entropies (using *m* = 3) of three hypothetical time series before (top row) and after (bottom row) the application of (a-b) linear detrending, (c-d) log-transformation, (e-f) interpolation of a missing or removed data point with a cubic smoothing spline. As these examples illustrate, with the exception of affine transformations, every pre-processing method discussed has the ability to alter our estimation of how much predictive information is contained in a time series. As such, performing pre-processing of a time series before permutation entropy is determined is not recommended. If the question to be addressed depends on such pre-processing, then care must be taken to understand how preprocessing is affecting the information estimate.

| Data processing method | Examples | Effect on |  | Remark |
| --- | --- | --- | --- | --- |
|  |  | Permutation entropy (PE) | Weighted permutation entropy (WPE) |  |
| Detrending | Linear, nonlinear (e.g., GAM), differencing | Increase or decrease | Increase or decrease | Effect will depend on attributes of the time series for any chosen permutation order $> 2$. |
| Transformation | $(x - \bar{x})/\sigma$, $\log(x)$, $\sqrt[4]{x}$, Fisher, etc. | None | Increase, decrease, or none | Normalization or rescaling will have no effect as long as the transformation is linear. Nonlinear transformations that compress large values (e.g., $\log(x)$) will increase WPE. Nonlinear transformations that amplify large values (e.g., Fisher) will decrease WPE. |
| Gap or sequence removal | Missing data (NAs), below detection level (zeros), species absences (zeros), constant values (poor precision) | Increase or decrease | Increase or decrease | Zeros should be retained if they represent true species absences (decreasing PE and WPE). Otherwise zeros and constant values can be removed (increasing or decreasing PE and WPE, **see main text**) or replaced by uncorrelated noise (increasing PE and WPE). The effect of concatenation will depend on attributes of the time series and gap size. Better to not count words that bridge gaps. |
| Interpolation | To infer gaps or to make time series equidistant | Increase or decrease | Increase or decrease | More likely to decrease than increase. Increases may occur for some nonlinear methods depending on attributes of the time series and the chosen permutation length. Better to ignore time-step uncertainty, assume equidistance, and not count words that bridge gaps. |
| Smoothing | Time-averaging,<br>time-summation | Decrease | Decrease | Like linear interpolation decreases PE and WPE by increasing the count of only-ascending or only-descending permutations. |
| Downsampling | Regular subsetting<br>to increase time-step<br>size | Increase or<br>decrease | Increase or<br>decrease | Effect will depend on attributes of the time series (particularly its intrinsic embedding dimension) and the chosen permutation length. |
| Time series<br>aggregation | Combining species<br>to functional group | Increase or<br>decrease | Increase or<br>decrease | Effect will depend on attributes of the time series being aggregated (e.g., their relative magnitudes, covariance, etc.). |

**Figure B2.**
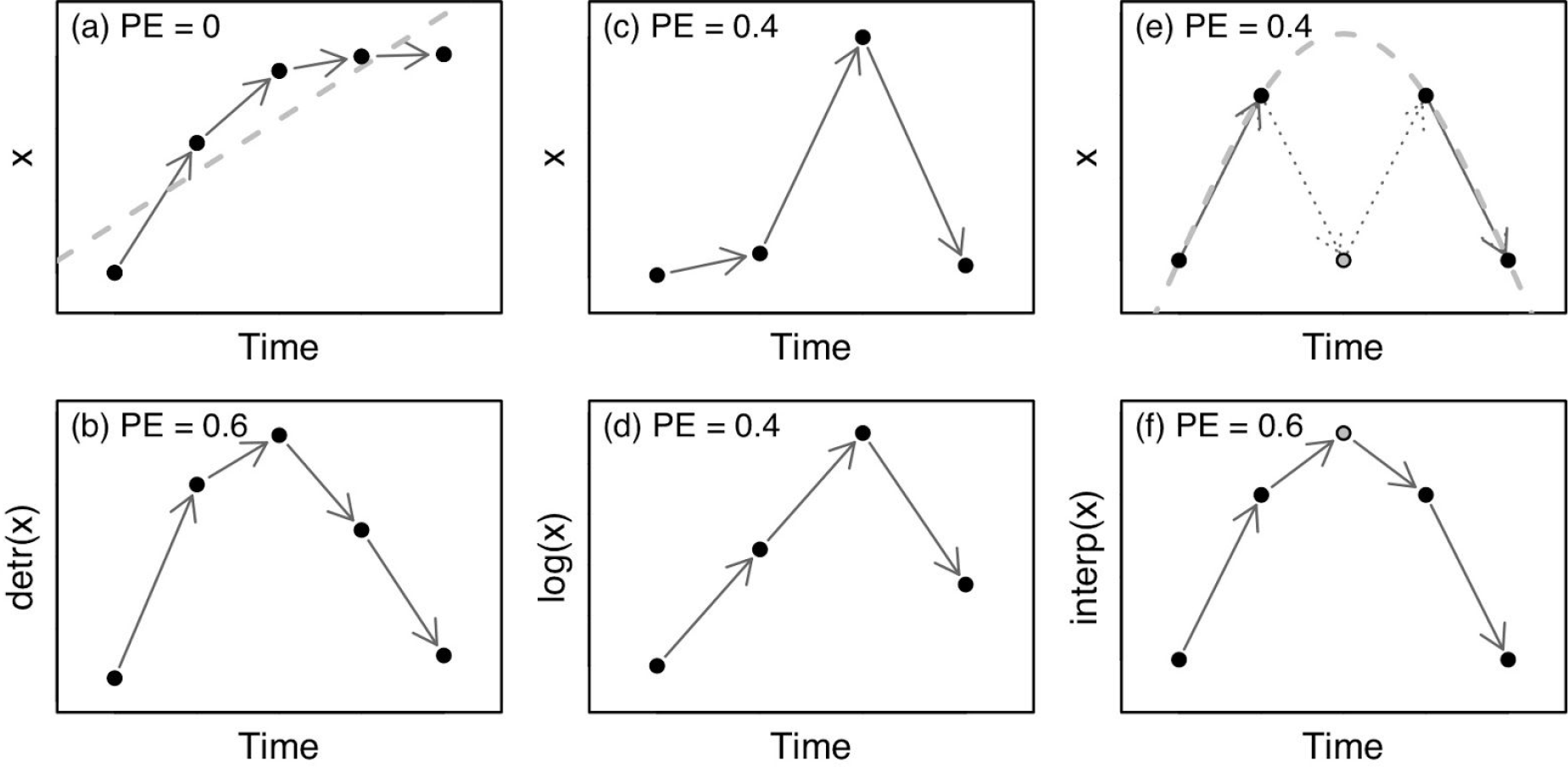
Anticipated effects of a suite of commonly used pre-processing methods on (non-weighted) permutation entropy (PE) using three hypothetical time series before (top row) and after (bottom row) the application of (a-b) linear detrending, (c-d) log-transformation, and (e-f) interpolation of a missing or removed data point with a cubic smoothing spline.

## Materials & Methods

### Forecasting with EDM

Empirical dynamic modelling is a set of nonlinear forecasting techniques brought to the attention of ecologists through the work of Sugihara (1994). The method is based on the idea that a system’s attractor generating the dynamics of a time series can be reconstructed via delay coordinate embedding (Takens 1981, Sauer et al. 1991), which can then be used to forecast system dynamics (e.g., Lorenz 1969, Weigend & Gershenfeld 1993, Farmer & Sidorowich 1987, Garland et al. 2015, Sauer in Weigend & Gershenfeld 1993, Casdagli & Eubank 1992, Smith 1992). The variant of these methods we use in this manuscript is based on the simplex projection and S-map method (Sugihara 1994) through the rEDM package.

The EDM approach first identifies the optimal embedding dimension *E* of the training data by fitting a model using simplex projection (Sugihara 1994). The embedding dimension *E* determines the number of temporal lags used for the delay coordinate embedding. We tested values for *E* between 1 and 10 and selected the value of *E* with the highest forecast skill using leave-one-out cross validation (Sugihara 1994). We then fitted the tuning parameter on the training data using the S-map model, describes the nonlinearity of the system and was varied in 18 steps between 0 and 10 (with log scaled intervals) to find the lowest error using leave-one-out cross validation on the training data.

In contrast to other forecasting methods such as ARIMA, the EDM approach searches across multiple time series models by finding the optimal in sample combination of embedding dimension and tuning parameter using cross-validation. Due to this model selection step, EDM tests a suite of forecasting models equal to the combination of and *E*. When is 0, the EDM model corresponds to an autoregressive model of the order of the embedding dimension (i.e. an AR3 model if E = 3). Values of **θ** greater than 0 can account for increasing degrees of state-dependence.

### Assessment of forecast error

We quantified forecasting error with the root mean squared error (Hyndman & Koehler 2006),

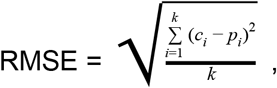

where *k* is the number of observed *c*_*i*_, values (i.e. abundances) and *p*_*i*_, are their corresponding predicted values. To compare forecast errors across time series that vary widely in units and variability, we normalized their RMSE by the range of observed values using

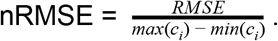

Smaller nRMSE corresponds to smaller forecasting error. Visual summaries of the observed and predicted dynamics are provided in the supplementary materials, including profiles of *E* and and the values chosen for predicting the test data.

### Calculation of permutation entropy

We calculated the weighted permutation entropy (WPE) of time series using the methods outlined in Box 1 (R code will be made available upon publication).

## Logistic Map Time Series

To demonstrate how both intrinsic and realized predictability change along a continuum from simple to complex and chaotic time series, we applied permutation entropy to time series from a well known population dynamic model, the Logistic Map:

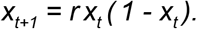

This model maps the current year’s population size to next year’s population size with simple density-dependence between non-overlapping generations. Although simple, this first-order, nonlinear function produces a wide range of dynamical behavior, from stable and oscillatory equilibria to chaotic dynamics (May 1976). We include this range of behavior by simulating the logistic map for 500 incremental growth rates between *r* = 3.4 and *r* = 3.9. We simulated each growth rate for 10,000 time steps keeping the last 3000 times steps for analysis. Weighted permutation entropy of time series was calculated for permutation order, *m*, from 3 to 5 and for time delay, τ, from 1 to 4. For simplicity, we will refer to weighted permutation entropy only in the results section and use the generic term permutation entropy everywhere else. Forecasting error for each time series was calculated using the normalized root mean squared error of an EDM forecast of the last 200 time steps.

Because ecological systems are influenced by both deterministic and stochastic drivers and the logistic map is purely deterministic, we sought to evaluate how stochasticity (noise) affects weighted permutation entropy and forecast error. To do so, we independently added both observational noise and process noise to the simulated population sizes by drawing random values from Gaussian distributions with standard deviations of either 0, 0.0001, 0.001, or 0.01 (Bandt & Pompe 2002). If the new population size was not between 0 and 1, a new value was drawn. Observational noise was added to the population size time series after the simulation, whereas process noise was added to population size at each time step during the simulation.

## Empirical Time Series Data

For empirical evidence of a relationship between permutation entropy and forecasting error, we examined a large variety of ecological time series that differ widely in complexity and data quality. We further investigated whether deviations from the expected general relationship can be explained by time series covariates such as measurement error (proxied here by whether the data originated from field versus lab studies), the nonlinearity of the time series (as quantified by the theta parameter of an EDM), or time series length. This allowed
us to identify possible predictors of time series complexity and the potential with which the time series of a system can be moved along the permutation-forecasting error (PE-FE) relationship to maximize realized predictability.

### Time series databases and processing

We compiled laboratory time series from the literature and field time series from the publicly available Global Population Dynamic Database (GPDD) for our analysis. The GPDD is the largest compilation of univariate time series available, spanning a wide range of geographic locations, biotopes and taxa (NERC Centre for Population Biology, 1999, Inchausti & Halley 2001). The GDPP database was accessed via the rGDPP package in R (https://github.com/ropensci/rgpdd). We added laboratory time series from studies by Becks et al. 2005, Fussmann et al. 2000, and the datasets used in a meta-analysis by Hiltunen et al. 2014. Time series with less than 30 observations, gaps greater than 1 time step and more than 15% of values being equal (and hence having the same rank in the ordinal analysis, i.e. ties) were excluded, resulting in a total of 461 time series. Each time series was divided into training (initial ⅔ of the time series) and test data (the last ⅓ of the time series), with the EDM model performing best on the training set being used to estimate forecast error in the test set. We calculated the weighted permutation entropy (WPE) of each empirical time series using a permutation order, *m*, of 3 and a time delay, **τ**, of 1. Results were robust to the choice of *m* ∈ [2, 5] and **τ** ∈ [1, 4], The three different ways to deal with ties (i.e. “random”, “first”, “average”) did not qualitatively affect the results, with results being robust to variation in time series minimum length and tie percentage

### Statistical analysis

All analyses were performed in the statistical computing environment R (R Development Core Team, 2018). We used the lme4 package to fit mixed models to investigate the relationship between forecast error and permutation entropy (Bates et al. 2015), with forecasting error being the dependent variable. The independent variables were permutation entropy, the data type, the number of observations (*N*), the proportion of zeros in the time series (zero_prop), the proportion of ties in the time series (ties_prop), and, from the EDM analysis, the nonlinearity ( ) and the embedding dimension (*E*) of the time series. The data type, i.e. whether time series were measured in the lab or in the field, was included with our hypothesis being that lab measurements have lower observation error. Zero and tie proportions were included as they pose problems to the estimation of PE, as do short time series (see Box 2).

## Results

### Logistic Map Time Series

The expected relationship between weighted permutation entropy and forecasting error occured in the simulations of the logistic map. Both WPE and FE generally increase as the growth rate, *r*, increases and the dynamics of the logistic map enter the realm of deterministic chaos (Fig. 2D). Correspondingly, both WPE and FE decline when chaotic dynamics converge to limit cycles (Fig. 2, gold example with r ≈ 3.84).

**Figure 2.**
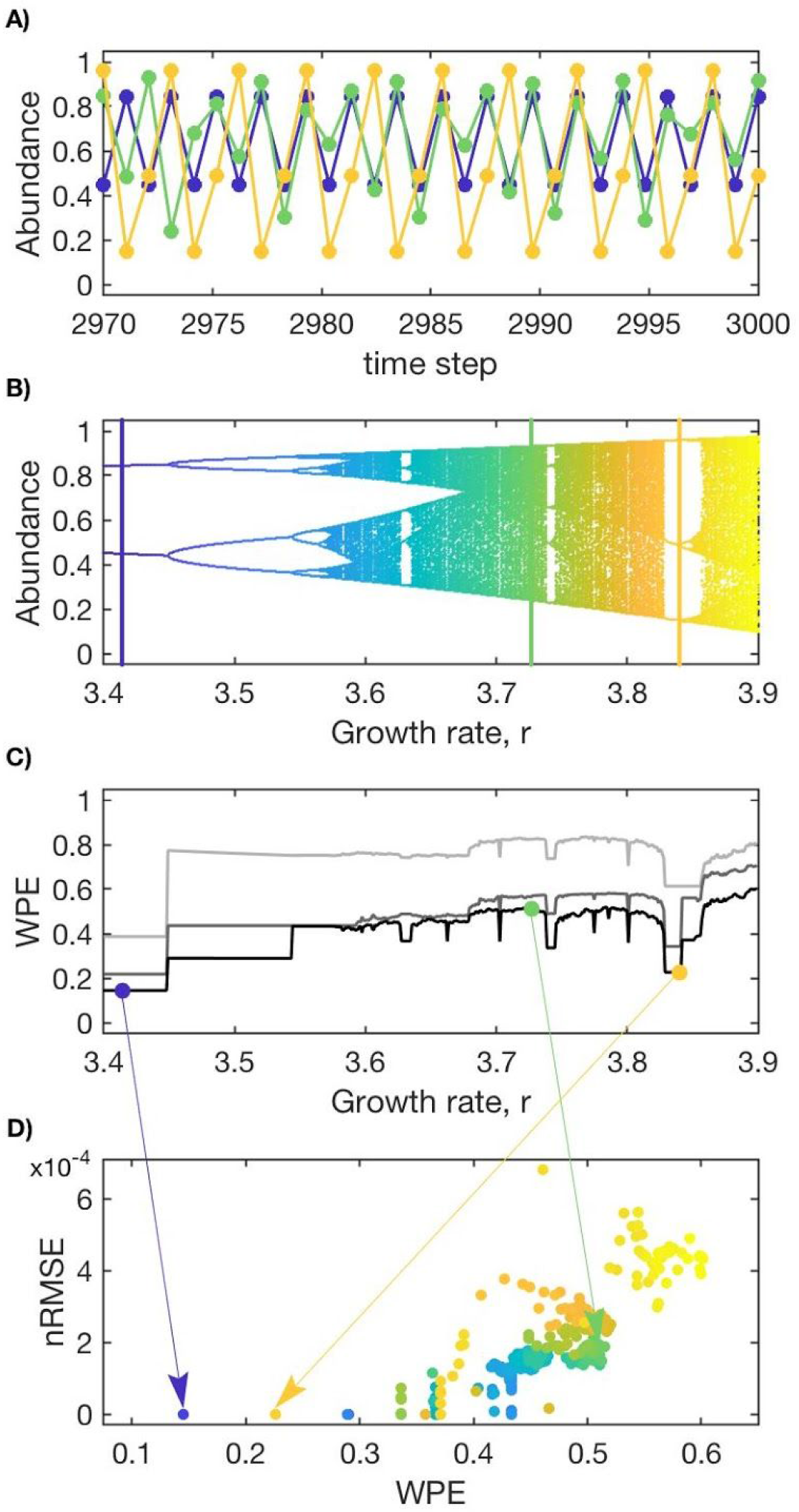
Simulations of the deterministic logistic map with no added process or observation noise. A) The last 30 time steps of three times series are plotted to demonstrate different behaviors, including 2-point limit cycles (r ≈ 3.41; dark blue), chaotic behavior (r ≈ 3.73; green), and 3-point limit cycles within the chaotic realm (r ≈ 3.84; gold). B) A bifurcation diagram of the logistic map attractor for growth rates between r = 3.4 and 3.9. C) Weighted permutation entropy (WPE) of the logistic map time series as the growth rate, *r*, changes for permutation order, *m*, of 3 (light grey), 4 (dark grey) and 5 (black), and time delay, of 1. D) forecast error quantified by the normalized root mean squared error (nRMSE) of an EDM forecast (*E* = 2, = 1) of the last 200 time steps of each simulation plotted against WPE (*m*=5, =1). The color coding corresponds to the growth rates in ‘B’.

The effect of stochastic noise on the WPE-FE relationship depended on the type of noise considered. While process noise strongly affects both WPE and FE (Fig. 3A) observational noise affects forecasting error more strongly than WPE (Fig. 3B). Indeed, the relationship between WPE and FE is largely obscured at high process noise but remains positive at high observational noise (Fig. 3A,B, top panels), particularly when dynamics are chaotic. When the dynamics are chaotic, the weighting in WPE is very effective at reducing the influence of observational noise on estimates of permutation entropy. However, when the dynamics exhibit stable limit cycles, WPE is sensitive to noise and this depends strongly on the chosen time delay, **τ**, and word length, *m*. For example, applying **τ**=2 for a 2-point limit cycle with a small amount of noise produces a WPE close to one, appearing as white noise as all permutations occur with equal probability. Limit cycles are best analyzed with **τ**=1 to capture the oscillations, although with *m*=3 small amounts of noise still result in two permutations occurring with equal frequency (1-3-2 or 2-3-1) and so WPE is elevated with respect to the no-noise case despite the high redundancy of the limit cycles (Fig. 3B, dark blue and gold points).

**Figure 3.**
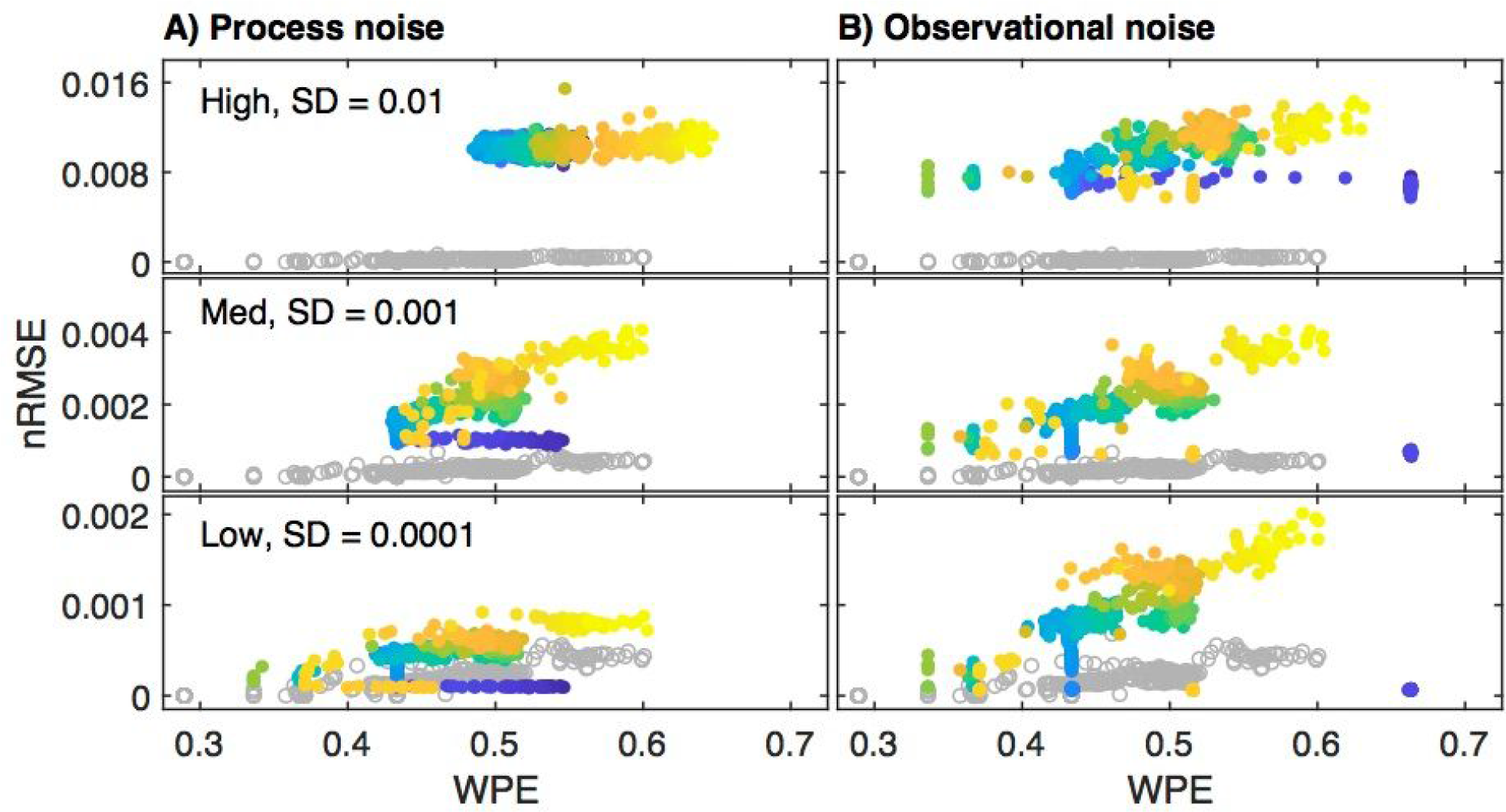
The relationship between weighted permutation error (WPE; *m*=5, =1) and forecasting error (measured as nRMSE) at three levels of A) process noise and B) observational noise. As the y-axis range and scale changes between subplots, the ‘no noise’ case is plotted in grey as a visual reference. The color coding corresponds to the growth rates in Fig. 2B. Systems with higher process noise exhibit both higher WPE and higher forecasting error. WPE is robust to observational noise when dynamics are chaotic, however limit cycles cause elevated estimates of WPE dependent on the choice of *m* and .

### Empirical Time Series Results

The 461 empirical time series vary in length (median = 50, min = 30, max = 197) and, as measured by WPE, complexity (median = 0.84, min = 0.076, max = 1). Forecasting error (nRMSE) ranges from 0.0000093 to 1.37, with a median of 0.19. Our analysis shows the expected positive relationship between permutation entropy and forecast error, with more complex time series (high WPE) yielding higher forecasting error (Table 1, center panel of Fig. 4). No difference in mean forecast error nor a difference in slope is detected between time series originating from lab or field studies (Table 1). Exploring the effects of time series covariates indicates that longer time series had lower FE, whereas time series with larger dimensionality (*E*) and greater nonlinearity (*θ*) as measured by EDM show higher FE (Table 1). These covariates increase the amount of variation in FE explained across time series to 35% (CI: 29 - 42%). An analysis of the partial R^2^ of all fixed effects in the model revealed that PE individually explained the largest amount of variation among predictors (21%, CI: 15 - 27%), followed by time series length (18%, CI: 12 - 24%), time series nonlinearity *θ* (6%, CI: 2 - 10%) and the chosen embedding dimension *E* (4%, 1 - 9%). Zero and tie proportions, as well as whether time series were from the field or the lab (type) explained less than 1% of the observed variation.

**Table 1:**
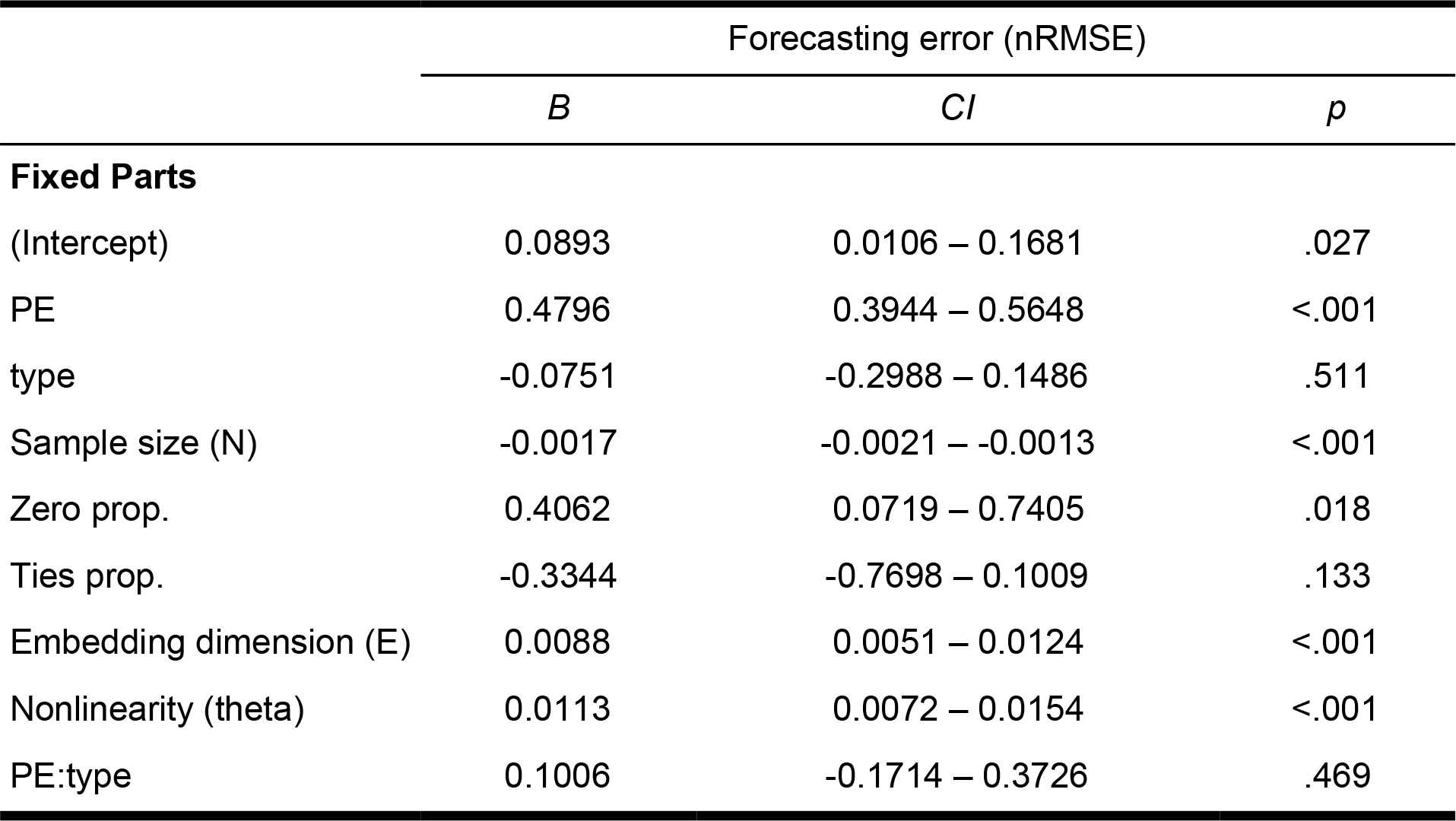
Model table presenting fixed effects of the mixed model analysis relating forecasting error to permutation entropy (PE), and additional time series covariates. Parameter estimates (*B*), 95% confidence intervals (CI) and p-values are provided. Forecasting error increases with weighted permutation entropy across 461 ecological time series

**Figure 4.**
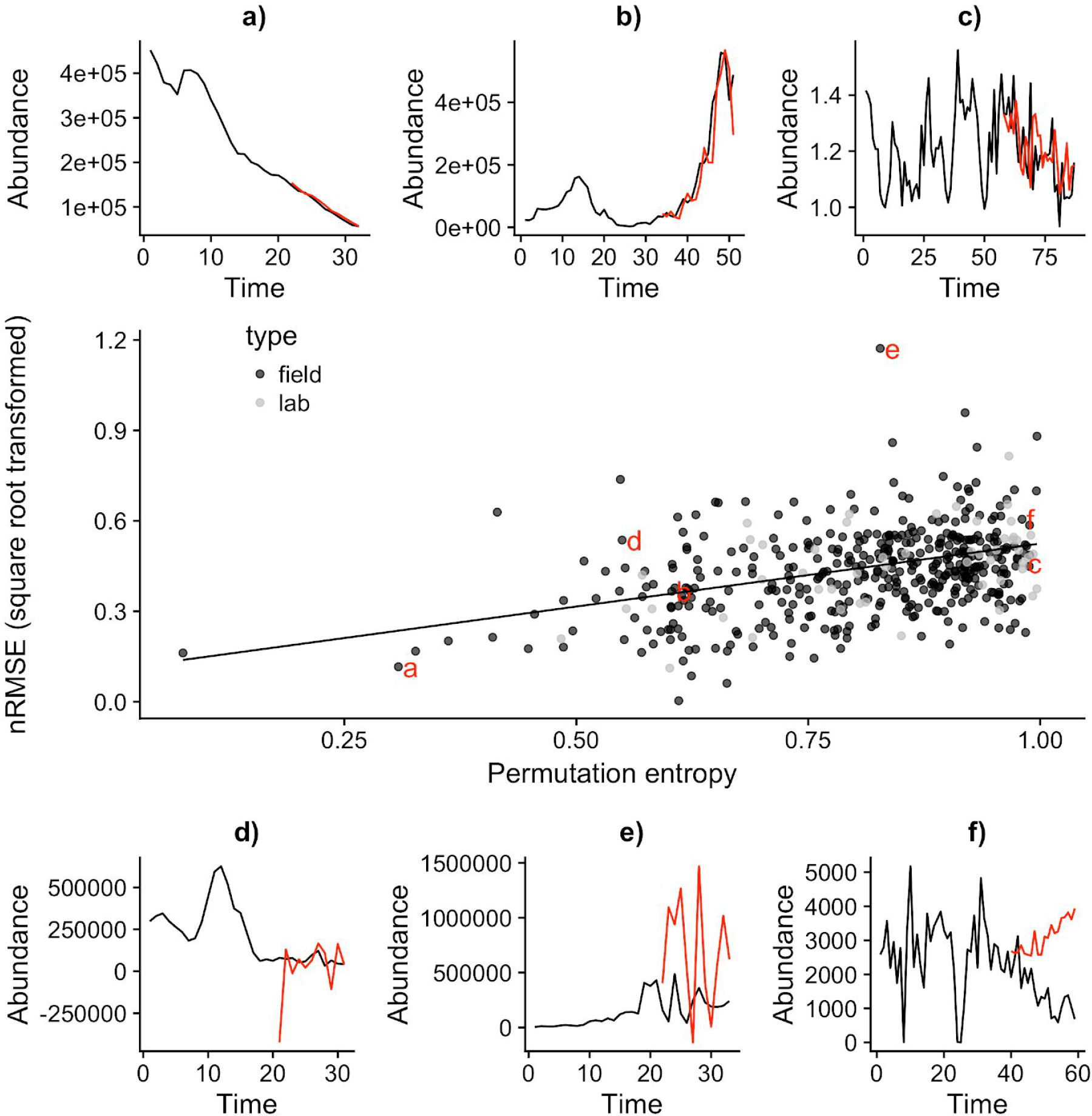
Relationship between weighted permutation entropy and forecast error (nRMSE, note square root scale of y axis) across 461 time series (middle panel) and specific exemplary time series (observations in black, forecasts in red, a-f). Forecast error increases with complexity of the time series as indicated by the higher permutation entropy value. The slope of the relationship was the same for time series from field and laboratory systems. The upper panels (a-c) show time series with forecast error lower than (a) or as expected (b-c) given their level of complexity, whereas the lower panels (d-f) illustrate time series which have higher than expected forecast error.

The PE v. FE relationship allows us to identify time series which were predicted better, equal to or worse than expected regarding their complexity (Fig. 4 a-f). Time series ‘b’ and ‘c’ fall along the expected relationship and hence are well predicted despite large differences in complexity. Time series ‘a’ shows a CIear trend which is well predicted. In contrast, time series ‘d’-‘f’ have higher than expected forecast error. Time series ‘d’ shows higher than expected error due to a strong outlier in the predicted values early in the test dataset. Time series ‘e’ is consistently poorly predicted, potentially due to wrong model choice or due to the short time series length. Time series ‘f’ is complex (high PE) with predictions missing the ongoing downward trend in the test data.

## Discussion

The urgent need for ecologists to provide operational forecasts to managers and decision makers requires that we understand when and why forecasts succeed or fail (Clark et al. 2001, Dietze 2017). We propose that the measurement of the intrinsic predictability of an ecological system can help reveal the origin of predictive uncertainty and indicate whether and how it can be reduced.

Our results show that realized and intrinsic predictability positively covary. The simulation study revealed that the relationship can be obscured by stochastic process noise, while measurement noise led to more scatter but preserved the positive slope (Fig 3). Although process noise often dominates over measurement noise in ecological time series (Ahrestani et al. 2013), the positive relationship between intrinsic and realized predictability we revealed across a wide range of empirical time series supports the applicability of our framework. In our analysis, permutation entropy explained the largest amount of variation (21%) in forecast error, followed by time series length, dimensionality and nonlinearity, jointly accounting for 35% of the variation. Time series that fell onto the expected relationship (Fig 4b,c) were well predicted given their complexity, whereas clear outliers (e.g. Fig 4e) would not require the use of PE to be identified as such. The relationship however allowed us to identify potential problems with forecasts of time series that have reasonable forecasts error, but which may be affected by overfitting (Fig 4a), outliers (Fig 4d) or regime shifts (Fig 4e) that may have gone unnoticed when looking at FE alone, particularly if applying automated or semi-automated forecasting methods across hundreds or thousands of time series (White et al. 2018).

### The value of intrinsic predictability to guide forecasting

A major advantage of permutation entropy is the independence from any assumed underlying model of the system, which makes this “model-free” method highly complementary to existing model-based approaches. For instance, Dietze (2017) recently proposed a model-based framework that partitions the contribution of various factors to predictive uncertainty, including the influence of initial conditions, internal dynamics, external forcing, parameter uncertainty and process error at different scales. If, for example, the dominant factor affecting near-term forecasts is deemed to be internal dynamics, then insight into intrinsic predictability would demonstrate how stable those internal dynamics are. Similarly, if a lot of variation remains unexplained by the model (i.e. the process error not explained by the known internal dynamics, initial conditions, external drivers, and estimated parameters), then “model-free” methods can provide insight into whether that variation is largely stochastic or contains unexploited structure that could be captured with further research into the driving deterministic processes. Finally, permutation entropy could be applied to the predicted dynamics of models to ascertain whether they accurately reflect properties of the observed dynamics, such as their complexity, similar to comparing the nonlinearity of a time series with the dynamics of the best model using the the EDM framework (Storch et al. 2017). Thus, intrinsic predictability provides diagnostic insights into predictive uncertainty and guidance for improving predictions.

### Comparative assessments of intrinsic predictability

The model-free nature of permutation entropy is advantageous in cross-system and cross-scale comparative studies of predictability. Whereas comparing all available forecasting methods on a given time series and predicting with the best-performing method would give us the best realized predictability (e.g. Ward et al. 2014), we would miss out on the comparative insight gained from aligning very different time series along the complexity gradient quantified by permutation entropy. Such a comparison could afford insight into whether intrinsic predictability differs across levels of ecological organization, taxonomic groups, habitats, geographic regions or anthropogenic impacts (Petchey et al. 2015). Determining the most appropriate covariates of monitored species (e.g. body size, life history traits, and trophic position) that minimize lost information would also inform monitoring methods. Furthermore, monitoring how realized and intrinsic predictability converge over time provides a means to judge improvements in predictive proficiency (Petchey et al. 2015, Houlahan et al. 2017, Dietze 2017). To do so, we need to apply available forecasting models to the same time series and measure their forecast error in combination with their intrinsic predictability. The monitoring of predictive proficiency has greatly advanced weather forecasting as a predictive science (Bauer et al. 2015). The analysis of univariate time series presented here only begins to put the intrinsic predictability of different systems into perspective. A primary goal is hence to expand the availability of long-term, highly resolved time series to determine potential covariates and improve our general understanding of ecological predictability (Petchey et al. 2015, Ward et al. 2014).

### Reliable assessment of intrinsic predictability

Permutation entropy requires time series data of suitable length and sampling frequency to infer the correct permutation order and time delay (Riedl et al. 2013). Long ecological time series measured at the appropriate time scales are rare, despite the knowledge that they are among the most effective approaches at resolving long-standing questions regarding environmental drivers (Lindenmeyer et al. 2012, Giron-Nava et al. 2017, Hughes et al. 2017). This problem is beginning to be resolved with automated measurements of system states, such as chlorophyll-a concentrations in aquatic systems (Thomas et al. 2018, Blauw et al. 2018), assessment of community dynamics in microbiology (Martin-Platero et al. 2018, Faust et al. 2015, Trosvik et al. 2008), and phenological (Pau et al. 2011) and flux measurements (Dietze 2017). Such high-frequency, long-term data are likely to provide a more accurate picture of the range of possible system states, even when systems are non-ergodic and change through time (e.g. Fig 4f). In fact, given the ease with which it is computed, PE can be assessed with a moving window across time or space to determine if a system is stationary or changing. As such, PE may be used as an early warning signal for system tipping points and critical transitions (Scheffer et al. 2009, Dakos et al. 2017) or to evaluate the effect of a management intervention on the system state.

Although the full potential of permutation entropy to guide forecasting is not yet realized, many other fields are starting to take advantage of its diagnostic potential. In paleoclimate science, permutation entropy has proven useful for detecting hidden data problems caused by outdated laboratory equipment (Garland et al. 2016), and in the environmental sciences it has provided insight into model-data deviations of gross primary productivity to further understand the global carbon cycle (Sippel et al. 2016). In epidemiology a recent study on the information-theoretic limits to forecasting of infectious diseases concluded that for most diseases the forecast horizon is often well beyond the time scale of outbreaks, implying prediction is likely to succeed (Scarpino & Petri 2017).

Our result showing that permutation entropy covaries with forecast error highlights the potential of using permutation entropy to better understand time series predictability in ecology and other disciplines.

## Acknowledgements

This paper originates from the “sPRED - Synthesizing Predictability Research of Ecological Dynamics” working group, supported by the Synthesis Centre of the German Centre for Integrative Biodiversity Research (DFG-FZT-118). FP, AT and OP benefitted from funding by the Swiss National Science Foundation (grant 31003A_159498 to OP). AI was supported by the Alexander von Humboldt Foundation. JG was supported by an Omidyar Fellowship from the Santa Fe Institute. HY is supported by the Gordon and Betty Moore Foundation’s Data-Driven Discovery Initiative through grant GBMF4563 to Ethan P. White. BR, BCR and UB acknowledge support by the German Research Foundation (FZT 118). We thank Gregor Fussmann and Lutz Becks for generously sharing time series data from microcosm experiments.

